# Underground gibberellin activity: differential gibberellin response in tomato shoots and roots

**DOI:** 10.1101/2020.07.27.222356

**Authors:** Uria Ramon, David Weiss, Natanella Illouz-Eliaz

## Abstract

Gibberellins (GAs) are growth-promoting hormones that regulate organ growth, mainly via cell elongation. Contradicting reports leave an open question of whether GA is as important for root elongation as it is for stem elongation.

Here we have addressed this question focusing on tomato (*Solanum lycopersicum*) primary-root elongation. We used a combination of physiological, molecular and genetic approaches to tackle this question.

Tomato has three GA receptors; GID1a, GID1b1 and GID1b2. The loss of all three receptors, strongly suppressed stem elongation and leaf expansion, but had a relatively minor effect on primary root elongation. The effect of GA on cell elongation and gene-expression was much weaker in roots, than in shoots, reaching saturation at lower hormone concentrations. Our results imply that this differential response to GA in shoots and roots is caused by the lower expression of the dominant GA receptor GID1a in roots.

We show that the differential activity of GA between shoots and roots affects root-to-shoot ratio, and speculate that this evolved as an adaptive mechanism to changing environments.

---

Plant organ growth is governed and modified by developmental programs and environmental cues. In most cases, these changes are mediated by the activity of phytohormones (Bradford and Trewaves, 1994; Verma *et al*., 2016). Gibberellins (GAs) are growth promoting hormones that regulate many developmental processes, including organ growth and elongation (Davière and Achard., 2013). GA affects elongation by promoting cell division and expansion (Ubeda-Thomas *et al*., 2009). The nuclear DELLA proteins inhibit all GA-elongating responses (Locascio *et al*., 2013) and GA binding to the GIBBERELLIN-INSENSITIVE DWARF1 (GID1) receptor leads to DELLA degradation and activation of growth (Ueguchi-Tanaka *et al*., 2005; Ueguchi-Tanaka *et al*., 2007).

While GA plays a central role in stem elongation (Sun and Gubler, 2004), its general significance for root elongation is less clear, with numerous conflicting reports (Torrey, 1976; Feldman, 1984; Phinney, 1984; Tanimoto, 2005; Tanimoto and Hirano, 2013). It is well established that Arabidopsis (*Arabidopsis thaliana*) root elongation is regulated by GA. Several studies demonstrate the central role of GA in Arabidopsis primary root elongation (Achard *et al*., 2009; Ubeda-Tomás *et al*., 2008; Ubeda-Tomás *et al*., 2009; Rizza *et al*., 2017). The Arabidopsis GA deficient mutant *ga1-3* exhibits shorter primary root, which is rescued by GA application or loss of DELLA activity (Fu and Harberd, 2003). Similarly, the loss of all three GA receptors strongly inhibit primary root elongation (Griffiths et al., 2016). Rizza *et al*., (2017) showed that endogenous bioactive GA levels correlate with cell length in Arabidopsis roots. Ubeda-Tomás *et al*. (2008) showed that inhibiting GA signaling specifically in the endodermis of Arabidopsis roots is sufficient to disrupt root elongation, indicating that the endodermis is the key site for GA action in the regulation of root elongation. This was supported by Shani *et al*. (2013) who found the accumulation of exogenous bioactive tagged-GAs in the endodermis of the elongation zone. Tanimoto and Hirano (2013) suggest that roots are very sensitive to GA and therefore respond to extremely low GA concentrations. For instance, while root elongation of the *ga1-3* Arabidopsis mutant was strongly induced by low concentration of GA_4_ (10^−10^ M), this treatment had no effect on leaf expansion (Arizumii *et al*., 2008).

In other plant species however, the role of GA in root elongation is unclear; while Whaley and Kephart (1957) show that GA application to maize (*Zea mays*) promotes root elongation, Svensson (1972) reported that the hormone has no effect on maize root growth. In Medicago (*Medicago truncatula*), GA treatment reduced, and GA-biosynthesis inhibitor paclobutrazol (PAC) increased, primary root length (Fonouni-Farde *et al*., 2019)., suggesting that GA inhibits root elongation, at least at higher concentrations. In tomato (*Solanum lycopersicum*), Butcher and Street (1960) show that elevating GA concentrations progressively promote root elongation, whereas Tognoni *et al*. (1966), showed that application of GA inhibits root elongation in a concentration-dependent manner. Barlow et al. (1991) demonstrate that roots of the GA-deficient mutant *gib-1*, grown *in vitro*, are shorter than WT, but this phenotype was not rescued by GA_3_ application. Thus, the role of GA in root elongation in species other than Arabidopsis, remains ambiguous.

To elucidate the role of GA in tomato primary-root elongation, we took advantage of the novel genetic resources, developed in our previous study (Illouz-Eliaz et al., 2019). We first examined primary root elongation in the GA deficient mutant *gib-2* (Koornneef *et al*., 1990). WT plants and *gib-2* mutants (both on cv. Moneymaker background) were grown in vermiculite in a growth room set to a photoperiod of 12/12-h night/days with light intensity (cool-white bulbs) of ∼250 μmol m^-2^ s^-1^ and 25°C. After two weeks, plants were repeatedly treated with 10^−5^ M GA_3_ and two-weeks later we measured stem and root length. Untreated *gib-2* stems were a third of the length of WT. GA application induced strong stem elongation in both genetic backgrounds, and their final length was similar (Fig. 1a). Non-treated *gib-2* primary roots, however, were not significantly shorter than those of the WT and GA treatment had only a very mild effect on root length (Fig. 1a). This observation suggests that either GA has no effect on tomato root elongation, or that root elongation is highly sensitive to GA and reaches saturation at very low levels of GA. Since *gib-2* exhibits residual GA activity (Illouz-Eliaz *et al*., 2019), this may be sufficient to allow normal root growth. We therefore tested primary root elongation in the *gid1* triple (*gid1*^*TRI*^) mutant that lacks any GA activity due to the loss of all three GA receptors, GID1a, GID1b1 and GID1b2 (Illouz-Eliaz *et al*., 2019). We first measured the elongation rate of primary roots and hypocotyls following germination on MS plates. Hypocotyl elongation rate of *gid1*^*TRI*^ was 10-times lower than that of WT (both on cv. M82 background, Fig. 1b), but root elongation rate of the mutant was only three-times lower (Fig. 1c). After ten days, primary roots of *gid1*^*TRI*^ were ca. half of the length of the WT, whereas the mutant hypocotyls were a tenth of the length of the WT (Fig. 1c). Root-to-shoot ratio was ca. four times higher in *gid1*^*TRI*^ (Fig. 1d). Since the *gid1*^*TRI*^ exhibits very slow growth, we also examined mature plants of the same physiological age (similar number of leaves in *gid1*^*TRI*^ and WT) grown in a greenhouse under natural day-length conditions with light intensity of 700 to 1000 µmol m^-2^ s^-1^ and 18-29°C. Primary root length of *gid1*^*TRI*^ were half of the WT, whereas *gid1*^*TRI*^ stems were a tenth of the WT (Fig 1e). Thus, the lack of GA activity, strongly affects shoot development, but only partially affects primary root elongation. We cannot exclude the possibility that the inhibition of root elongation in *gid1*^*TRI*^ was a result of limited assimilate supply by the extremely small canopy.

**Figure 1.**
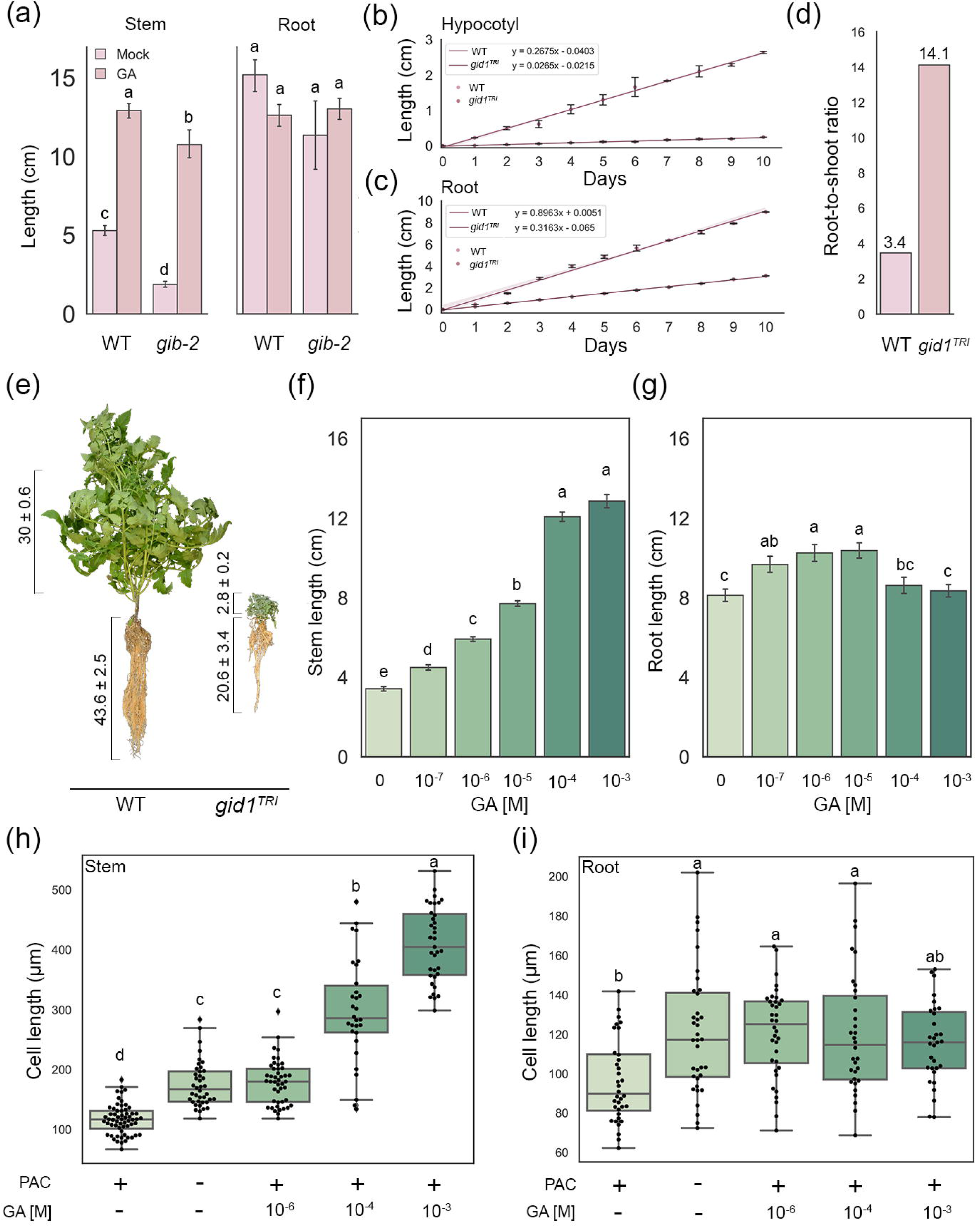
GA-induced elongation response spatially differentiates in tomato roots and shoots (**a**) Stem and root elongation of 14-days-old WT and *gib-2* mutants (cv. Moneymaker), treated continuously (or not) with 10^−5^ M GA_3._ Values are mean of 4 biological replicates ± SE. Each set of letters above the columns represents significant differences (Tukey–Kramer HSD, P < 0.05). (**b** and **c**) Hypocotyl (b) and root (c) elongation rate of WT and *gid1*^*TRI*^ (cv. M82) seedlings grown on MS plates. Values are mean of 7 seedlings of each line and the shadow presents the statistical mean. (**d**) Root-to-shoot ratio of 10-day-old WT and *gid1*^*TRI*^ seedlings. (**e**) Representative image of WT and *gid1*^*TRI*^ at the same physiological age. Numbers present the average length in cm of 3 biological replicates ± SE. (**f** and **g**) Stem (f) and primary root (g) length of 14-day-old WT (cv. M82) seedlings treated with 2 mg/l PAC followed by GA_3_ application (10^−7^ to 10^−3^M). Values are mean of 10 plants ± SE. Each set of letters above the columns represents significant differences (student’s t test, p < 0.05). (**h** and **i**) Epidermal cell length of elongating stems (h) and primary roots at the elongation zone (i), was measures after 10 days of treatment with PAC or PAC + GA_3_ using confocal microscopy and analyzed by imageJ. Each set of letters above the columns represents significant differences (student’s t test, p < 0.05).

We further tested the effect of exogenous GA on root and stem elongation. To this end, 14-day-old WT (cv. M82) seedlings grown in vermiculite as described above, were treated with 2 mg/l PAC followed by the application of elevating GA_3_ concentrations (from 10^−7^ M to 10^−3^ M). After 15 days, we measured stem and primary root length. WT stems exhibited a strong and increased elongation response to rising GA_3_ concentrations, reaching saturation at very high concentrations (above 10^−5^ M, Fig. 1f). In contrast, primary root length was hardly affected and exhibited a bell-curve response (Fig. 1g). To examine how GA affects cell length, we treated WT seedlings with PAC or PAC with GA_3_ (10^−6^, 10^−4^, and 10^−3^ M), and after 10 days we measured epicotyl and primary root epidermal cell length using confocal microscopy. Stem epidermal cell elongation strongly responded to GA treatments and cell length of stems treated with PAC and 10^−3^ M GA_3_ were four times longer than those treated only with PAC (Fig 1h). Primary root epidermal cells exhibited a very mild response to PAC and GA_3_ (Fig 1i) and the effect of the hormone was saturated already at 10^−6^ M. GA-treated cells were only 1.2 times longer than the PAC treated cells. It is worth noting that higher GA concentrations did not inhibit root cell length, suggesting that reduced cell number may be the cause for inhibition of root length in high GA concentration.

It was previously suggested that roots are more sensitive to GA than stems (Tanimoto and Hirano, 2013). To further test root sensitivity to GA, we analyzed the response of various known GA-regulated genes, including GA biosynthesis (*GA 20-OXIDASE*s and *GA 3-OXIDASE*) and signaling (*GID1s*) genes that are downregulated by the hormone (Middleton *et al*., 2012; Illouz-Eliaz *et al*., 2019) and the GA-induced gene *GIBBERELLIC ACID STIMULATED TRANSCRIPT1* (*GAST1*, Shi *et al*., 1992). Seedlings, grown on vermiculite as described above, were treated with 2 mg/l PAC for 10 days and then with several GA_3_ concentration (10^−8^ to 10^−5^ M), and three hours later, gene expression in elongating stems and roots was analyzed by qRT-PCR as described by Illouz-Eliaz et al. (2019). We found a clear and significant response to 10^−7^ M, but not to 10^−8^ M GA_3_, for all genes, in both elongating stems and roots (Fig 2a-f), suggesting similar sensitivity to GA. Moreover, the intensity of the molecular response in roots was weaker and saturated at lower concentrations than in stems. These results are consistent with our cell elongation results (Fig. 1h and i).

**Figure 2.**
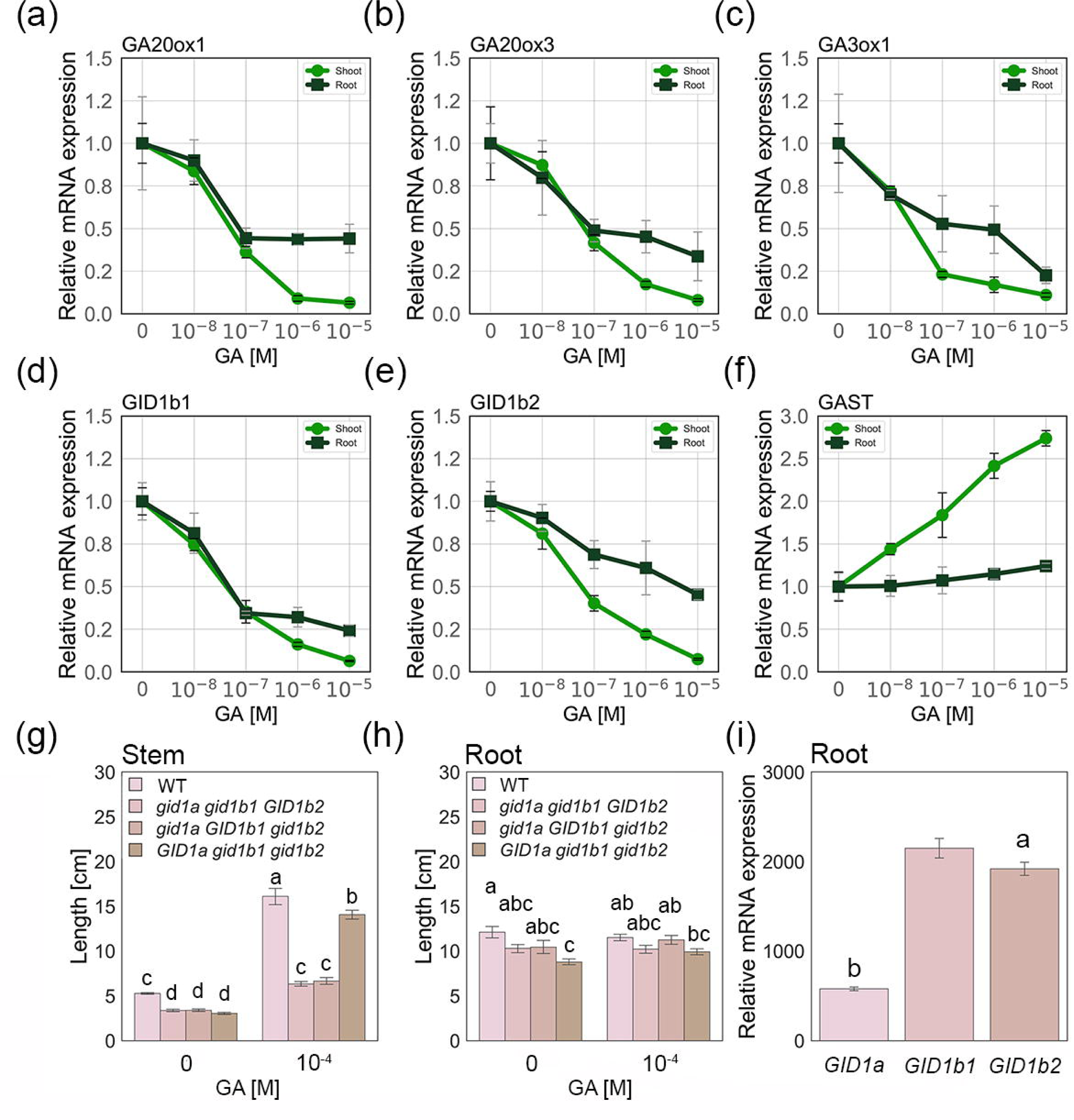
qRT-PCR analysis of (**a**) *GA 20-OXIDASE-1* (*GA20ox-1*) (**b**) *GA20ox-3* (**c**) *GA3ox-1* (**d**) *GID1b1* (**e**) *GID1b2* and (**f**) *GAST1* expression in elongating stems and roots treated with 2 mg/l PAC. After 10 days with PAC, plants were treated with different GA_3_ concentrations and RNA was extracted three hours later for expression analysis. Values (normalized to ACTIN) are means of four biological replicates ± SE. (**g** and **h**) Stem (g) and primary root (h) length of 14-day-old WT and *gid1* double mutant (cv. M82) seedlings treated with 2 mg/l PAC followed by GA_3_ application (10^−4^M). Values are mean of 9 plants ± SE. Each set of letters above the columns represents significant differences (Tukey–Kramer HSD, P < 0.05). (**i**) Tomato *GID1a, GID1b2* and *GID1b2* expression in roots taken from transcriptomic data by Góra-Sochacka *et al*., 2019. Values are mean of 3 plants ± SE. Each set of letters above the columns represents significant differences (student’s t test, p < 0.05).

We previously showed that GID1a is the dominant GA receptor in tomato stems, due to its high affinity to DELLA and the fact that it is not inhibited by the feedback response to GA (Illouz-Eliaz *et al*., 2019). Its presence alone, in the absence of GID1b1 and GID1b2 activity, can induce the strong stem-elongation response to exogenous GA that is saturated only at very high concentrations. GID1b1 and GID1b2 exhibit a weak GA response that is saturated at low concentrations. To examine if the differential response to GA in stems and roots results from differential activity of the different GID1s, we tested the effect of GA application on the three tomato *gid1* double mutants. The GID1 family in tomato is composed of three members, therefore each double mutant contains one active GID1. In stems, only *gid1b1 gid1b2* with active GID1a exhibited strong elongation response, similar to WT (Fig. 2g, Illouz-Eliaz *et al*., 2019). In contrast, roots of this double mutant did not show elongation response (Fig. 2h), suggesting that GID1a does not promote strong GA response in roots as it does in stems. Previously we showed that *GID1a* and *GID1b1* are highly expressed in elongating stems, while *GID1b2* exhibits relatively low expression (Illouz-Eliaz *et al*., 2019). We analyzed available public data of tomato root transcriptome (Koenig *et al*., 2013; Zouine *et al*., 2017; Góra-Sochacka *et al*., 2019; Gray *et al*., 2020) and found very low expression of *GID1a* compared to *GID1b1* and *GID1b2* in all datasets (Fig. 2i presents the data analyzed from Góra-Sochacka *et al*., 2019). This raises the possibility that the minor effect of GA on root elongation, is caused by low *GID1a* expression in roots. Further research regarding the activity of GID1a as a disjunctive component of GA signaling in above-and underground organs, is thus highly warranted. An example for such studies could be exploring the effect of highly expressed GID1a under a root-specific promoter.

To conclude, our results in tomato suggest that while GA plays a central role in the regulation of shoot elongation, it has only a minor effect on root elongation. Although previous studies suggest that roots are more sensitive to GA than shoots, this is probably not the case in tomato. We found however that very low GA concentrations are sufficient to saturate root elongation, but not stem elongation.

Plants adjust root-to-shoot ratio to adapt to changes in the environment. Under drought conditions root-to-shoot ratio increases to reduce transpiration and increase water uptake (Xu *et al*., 2015). GA accumulation is inhibited under osmotic stresses, such as drought and salinity (Achard *et al*., 2006; Colebrook *et al*., 2014). Thus, the reduced GA levels strongly affects shoot but not root growth. It is possible that this mechanism evolved as a strategy to modify root-to-shoot ratio under stress conditions.

## Acknowledgement

This research was supported by a research grants from the Israel Ministry of Agriculture and Rural Development (12-01-0014), the Israel Ministry of Agriculture and Rural Development (Eugene Kandel Knowledge Center) as part of the “Root of the Matter”-The Root zone knowledge center for leveraging modern agriculture, and by the Israel Science Foundation (grant No. 617/20) to D.W. We thank Prof. Naomi Ori, Dr. Idan Efroni and Prof. Eilon Shani for helpful discussion.

## References

Achard P, Cheng H, De Grauwe L, Decat J, Schoutteten H, Moritz T, Van Der Straeten D, Peng J, Harberd NP. 2006. Integration of plant responses to environmentally activated phytohormonal signals. Science 311: 91–94.

Achard P, Gusti A, Cheminant S, Alioua M, Dhondt S, Coppens F, Beemster GTS, Genschik P. 2009. Gibberellin signaling controls cell proliferation rate in Arabidopsis. Current Biology 19: 1188–1193.

Ariizumi T, Murase K, Sun T, Steber CM. 2008. Proteolysis-Independent Downregulation of DELLA repression in Arabidopsis by the gibberellin receptor GIBBERELLIN INSENSITIVE DWARF1. The Plant Cell 20: 2447–2459.

Barlow PW, Brain P, Parker JS. 1991. Cellular growth in roots of a gibberellin-deficient mutant of tomato (*Lycopersicon escolentum* Mill.) and its wild-type. Journal of Expiramental Botany 42: 339–351.

Bradford KJ, Trewavas AJ. 1994. Sensitivity thresholds and variable time scales in plant hormone action. Plant Physiology 105: 1029–1036.

Butcher DN, Street HE. 1960. Effects of kinetin on the growth of excised tomato roots. Physiologia Plantarum 13: 46–55.

Colebrook EH, Thomas SG, Phillips AL, Hedden P. 2014. The role of gibberellin signalling in plant responses to abiotic stress. Journal of Experimental Biology 217: 67–75.

Davière J, Achard P. 2008. Gibberellin signaling in plants. Development 140: 1147–1151.

Feldman LJ. 1984. Regulation of root development. Annual review of plant physiology 35: 223–242.

Fonouni-Farde C, Miassod A, Laffont C, Morin H, Bendahmane A, Diet A, Frugier F. 2019. Gibberellins negatively regulate the development of *Medicago truncatula* root system. Scientific Reports 9: 1–9.

Fu X, Harberd NP. 2003. Auxin promotes Arabidopsis root growth by modulating gibberellin response. Nature 421: 740–743.

Góra-Sochacka A, Wiesyk A, Fogtmann A, Lirski M, Zagórski-Ostoja W. 2019. Root transcriptomic analysis reveals global changes induced by systemic infection of solanum lycopersicum with mild and severe variants of potato spindle tuber viroid. Viruses 11: 992.

Gray SB, Rodriguez-Medina J, Rusoff S, Toal TW, Kajala K, Runcie DE, Brady SM. 2020. Translational regulation contributes to the elevated CO2 response in two Solanum species. The Plant Journal 102: 383–397.

Griffiths J, Murase K, Rieu I, Zentella R, Zhang Z, Powers SJ, Gong F, Phillips AL, Hedden P, Sun T, et al. 2004. Genetic characterization and functional analysis of the GID1 gibberellin receptors in Arabidopsis. The Plant Cell 18: 3399–3414.

Illouz-Eliaz N, Ramon U, Shohat H, Blum S, Livne S, Mendelson D, Weiss D. 2019. Multiple gibberellin receptors contribute to phenotypic stability under changing environments. The Plant Cell 31: 1506–1519.

Koenig D, Jiménez-Gómez JM, Kimura S, Fulop D, Chitwood DH, Headland LR, Kumar R, Covington MF, Devisetty UK, Tat AV, et al. 2013. Comparative transcriptomics reveals patterns of selection in domesticated and wild tomato. Proceedings of the National Academy of Sciences of the United States of America 110: 2–9.

Koornneef M, Bosma TDG, Hanhart CJ, Veen H Van Der, Zeevaart JAD. 1990. The isolation and characterization of gibberellin-deficient mutants in tomato. Theoretical and Applied Genetics 80: 852–857.

Locascio A, Blázquez MA, Alabadi D. 2013. Genomic analysis of DELLA protein activity. Plant and Cell Physiology 54: 1229–1237.

Middleton AM, Úbeda-tomás S, Griffiths J, Holman T, Hedden P, Thomas SG, Phillips AL, Holdsworth MJ, Bennett MJ, King JR, et al. 2012. Mathematical modeling elucidates the role of transcriptional feedback in gibberellin signaling. Proceedings of the National Academy of Sciences 109: 7571–7576.

Nakajima M, Shimada A, Takashi Y, Kim Y, Park S, Ueguchi- M, Suzuki H, Katoh E, Iuchi S, Kobayashi M, et al. 2006. Identification and characterization of Arabidopsis gibberellin receptors. Plant Journal 46: 880–889.

Phinney BO. 1985. Gibberellin A1 dwarfism and shoot elongation in higher plants. Biologia Plantarium 27: 172–179.

Rizza A, Walia A, Lanquar V, Frommer WB, Jones AM. 2017. In vivo gibberellin gradients visualized in rapidly elongating tissues. Nature Plants 3: 803–813.

Shani E, Weinstain R, Zhang Y, Castillejo C, Kaiserli E, Chory J, Tsien RY, Estelle M. 2013. Gibberellins accumulate in the elongating endodermal cells of Arabidopsis root. Proceedings of the National Academy of Sciences of the United States of America 110: 4834–4839.

Shi L, Gast RT, Gopalraj M, Olszewski NE. 1992. Characterization of a shoot-specific, GA- and ABA-regulated gene from tomato. Plant Journal 2: 153–159.

Sun T, Gubler F. 2004. Molecular mechanism of gibberellin signaling in plants. Annual Review of Plant Biology 55: 197–223.

Svensson SB. 1972. A comparative study of the changes in root growth, induced by coumarin, auxin, ethylene, kinetin and gibberellic acid. Physiologia Plantarum 26: 115–135.

Tanimoto E. 2005. Regulation of root growth by plant hormones-roles for auxin and gibberellin. Critical Reviews in Plant Sciences 24: 249–265.

Tanimoto E, Hirano K. 2013. Role of gibberellins in root growth. In: Eshel A, Beeckman T, eds. Plant roots: the hidden half. CRC Press 1–14.

Tognoni F, Halevy AH, Wittwer SH. 1967. Growth of bean and tomato plants as affected by root absorbed growth substances and atmospheric carbon dioxide. Planta 72: 43–52.

Torrey JG. 1976. Root hormones and plant growth. Annual Review of Plant Physiology 27: 435–459.

Ubeda-Tomás S, Federici F, Casimiro I, Beemster GTS, Bhalerao R, Swarup R, Doerner P, Haseloff J, Bennett MJ. 2009. Gibberellin signaling in the endodermis controls Arabidopsis root meristem size. Current Biology 19: 1194–1199.

Ubeda-Tomás S, Swarup R, Coates J, Swarup K, Laplaze L, Beemster GTS, Hedden P, Bhalerao R, Bennett MJ. 2008. Root growth in Arabidopsis requires gibberellin/DELLA signalling in the endodermis. Nature Cell Biology 10: 625–628.

Ueguchi-Tanaka M, Ashikari M, Nakajima M, Itoh H, Katoh E, Kobayashi M, Chow T, Hsing YC, Kitano H, Yamaguchi I, et al. 2005. GIBBERELLIN INSENSITIVE DWARF1 encodes a soluble receptor for gibberellin. Nature 437: 693–698.

Ueguchi-tanaka M, Nakajima M, Katoh E, Ohmiya H, Asano K, Saji S, Hongyu X, Ashikari M, Kitano H, Yamaguchi I, et al. 2007. Molecular interactions of a soluble gibberellin receptor, GID1, with a rice DELLA protein, SLR1, and gibberellin. The Plant Cell 19: 2140–2155.

Verma V, Ravindran P, Kumar PP. 2016. Plant hormone-mediated regulation of stress responses. BMC Plant Biology 16: 1–10.

Whaley WG, Kephart J. 1957. Effect of gibberellic acid on growth of maize roots. Science 125: 234.

Xu W, Cui K, Xu A, Nie L, Huang J, Peng S. 2015. Drought stress condition increases root to shoot ratio via alteration of carbohydrate partitioning and enzymatic activity in rice seedlings. Acta Physiologiae Plantarum 37: 9.

Yamamoto Y, Hirai T, Yamamoto E, Kawamura M, Sato T, Kitano H, Matsuoka M, Ueguchi-tanaka M. 2010. A Rice *gid1* suppressor mutant reveals that gibberellin is not always required for interaction between its receptor, GID1, and DELLA proteins. The Plant Cell 22: 3589–3602.

Zouine M, Maza E, Djari A, Lauvernier M, Frasse P, Smouni A, Pirrello J, Bouzayen M. 2017. TomExpress, a unified tomato RNA-Seq platform for visualization of expression data, clustering and correlation networks. Plant Journal 92: 727–735.

